# Single-Cell Modeling of CD8^+^ T Cell Exhaustion Predicts Response to Cancer Immunotherapy

**DOI:** 10.1101/459867

**Authors:** Guangxu Jin, Gang Xue, Rui-Sheng Wang, Ling-Yun Wu, Lance Miller, Yong Lu, Wei Zhang

## Abstract

Accurate prediction of response to immune checkpoint blockade (ICB) and simultaneous development of new efficacious ICB strategy are unmet needs for cancer immunotherapy^1-5^. Despite the advance of existing clinical or computational biomarkers^6-10^, it is very limited in explicating phenotypic variability of tumor-infiltrating CD8^+^ T lymphocytes (TILs) at the single-cell level, including tumor-specificity^11,12^ and distinct exhaustion profiles^13-16^, that are crucial for the responsiveness to ICB. Here we show that a quantitative criterion, D value, for evaluating tumor-specific TIL exhaustion, identified by our new high-dimensional single-cell-based computational method, called HD-scMed, accurately predicts response to the ICB, i.e., αPD-1, αCTLA-4, and αPD-1+αCTLA-4, with the performance of AUC=100% in human tumors. D-value is the Euclidean distance from a subset of exhausted CD8^+^ TILs, identified as Pareto Front (PF)^18-21^ by HD-scMed within the high-dimensional expression space of a variety of exhaustion markers, to the baseline TILs excluded by the PF. We phenotypically distinguished two types of TIL “exhaustion” by the D value, namely “D Extremely-high” in non-responders, associating with high tumor-specificity imprinted with enhanced exhaustion and inactivated effector and cytotoxic signatures; and “D low” specific to responders, alternatively enriched with T cell activation and cytolytic effector T cell signatures. We also observed a large portion of “D negative” bystander T cells irrelevant to response. Notably, D-low TILs in clinical responders display very low LAG3 expression. To reverse the functionality of the LAG3-high TILs after receiving αPD-1+αCTLA-4, we combined αLAG3 with αPD-1+αCTLA-4 in treating a murine tumor model. Remarkably, αLAG3+αPD-1+αCTLA-4 displays extraordinary antitumor efficacy to eradicate advanced tumors, which is associated with burst GZMB^hi^ TIL populations; whereas αPD-1+αCTLA-4 or αPD-1+αCTLA-4 plus other ICBs as control only induce moderate tumor growth inhibition. Our study has important implications for cancer immunotherapy as providing both accurate predictions of ICB response and alternative strategy to reverse ICB resistance.

## Main Text

Characteristics of TIL exhaustion are generally considered as low effector function, decreased proliferation, and diminished cytokine production^13-16^. However, the challenges in understanding the key role of TIL exhaustion, especially in determining responses to ICBs, come with the identification of appropriate exhaustion biomarker or criterion. At the single-cell level, the quantitative exhaustion criterion is required to be able to demonstrate fundamentally distinct types of T cell functions represented by various functional molecules, such as, surface exhaustion markers (PD-1, CTLA-4, LAG3, TIM3, VISTA, TIGIT, etc)^22-26^, related transcriptional factors (EOMES, T-BET, PRDM1, etc)^15^, effector molecules (IFNG, GZMB, GZMA, PRF1, FASLG)^27^, and tumor-specificity markers (CD39 and CD103)^11,12^. The developed D value as such a quantitative exhaustion criterion models TILs within a high-dimensional expression space defined by a number of exhaustion related markers (**Online Methods**, **Extended Data Fig. 1**, **Supplementary Methods**). To predict response to ICB, we used two single-cell mass cytometry (CyTOF) datasets^17^, including 11 specimens from 7 melanoma patients received the ICBs of αPD-1, αCTLA-4, or αPD-1+αCTLA-4 and another 34 mouse specimens from melanoma mouse tumors received the ICBs of αPD-1 or αCTLA-4 (**Online Methods**, **Extended Data Fig. 2**-**3**, **Supplementary Table 1-2**). D-values for the tumors were identified from a 11-dimensional expression space (immune checkpoints or ICs: PD-1, CTLA-4, LAG3, and TIM3, other relevant markers: KLRG1, Blimp1, BCL6, Ki-67, and CD127, and the related transcriptional factors or TFs: EOMES and T-bet). The proportion of the PF single cells in all TILs is 18%±5% (**Fig. 1a**, **Supplementary Table 3**). We identified that the exhausted CD8^+^ TILs of the non-responders have significantly higher D-values than those of responders (*P* < 10^−16^, Mann-Whitney *U* test), refer to **Fig. 1b,e**. The proportion of D-high TILs in non-responders is significantly higher than that in responders (*P* < 10^−15^, Fisher’s exact test, **Fig. 1c-d**). By analyzing the contributions (DE_i_) of the exhaustion markers to D values, we observed significant contributions from PD-1, CTLA-4, and LAG3 to high D values in non-responder (**Fig. 1g**, *P* < 10^−16^, Mann-Whitney *U* test). Thus, D value predicts clinical response to ICBs in human tumors accurately, that is, AUC=100% (**Online Methods**, **Fig. 1f**). Similarly, D value predicts ICB responses of mouse tumors at the accuracy of AUC=83% (**Extended Data Fig. 4c**). More importantly, the prediction accuracy is not affected by the number of exhaustion markers used in the model (**Extended Data Fig. 9**, an example with 6 markers).

**Figure 1.**
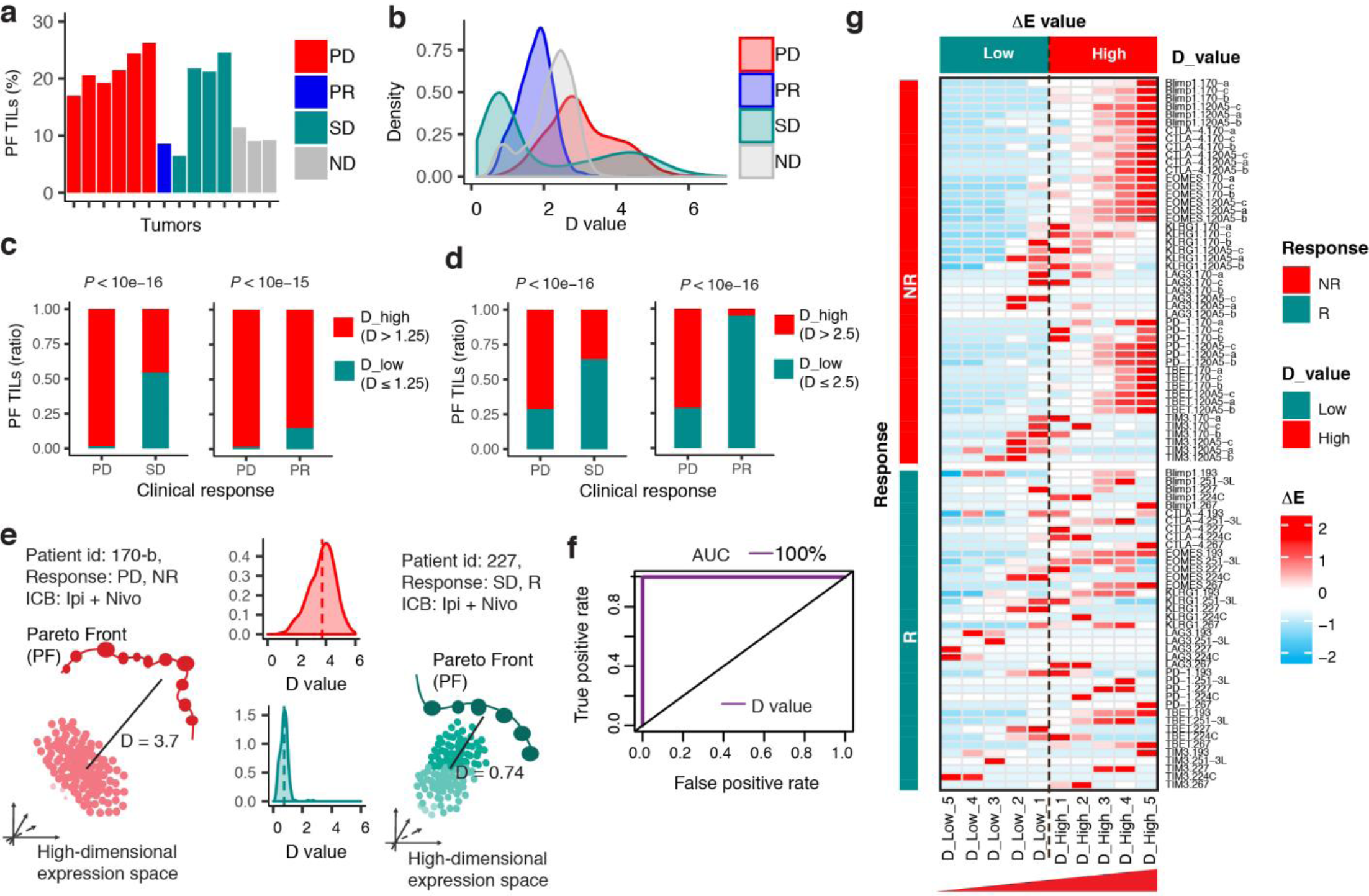
D value from HD-scMed in distinguishing clinical responses to ICB. Clinical responses include PD: progressive disease, PR: partial response, and SD: stable disease. ICBs are αPD-1, αCTLA-4, and αPD-1+αCTLA-4. ND: PBMC samples from healthy donors were used as the baseline for ICB samples. **(a).** Identification of **Pareto front (PF)** TILs as exhausted TILs in the high-dimensional expression space. TIL exhaustion is evaluated by HD-scMed in the high-dimensional expression space defined by PD-1, CTLA4, LAG3, EOMES, TIM3, and TBET, from human CyTOF antibody panel (**Online Methods**, **Extended Data Fig. 1**). Shown is a small portion of PF TILs identified as exhausted TILs by HD-scMed. **(b).** D value distributions in tumors with different clinical responses. D value is defined as the Euclidean distance or straight-line distance in the 11-dimensional expression space by the expression levels of the 11 exhaustion markers. D value for each PF TIL is the Euclidean distance between the PF TIL and its baseline non-front TILs that are dominated by the PF TIL (**Online Methods**, **Extended Data Fig. 1a-b**). **(c).** Analysis of D-low and D-high PF TILs of the tumors from non-responders (PD) versus responders (PR or SD) by the threshold of D = 1.25. Fisher’s exact test. **(d).** Analysis of D-low and D-high PF TILs of the tumors from non-responders (PD) versus responders (PR or SD) by the threshold of D = 2.5. Fisher’s exact test. **(e)**. An instance to show the difference of D-value in the TILs of a non-responder (NR) and a responder (R). Both patients received the combination therapy of αPD-1+αCTLA-4 by ipilimumab+nivolumab. Tumor 170-b is from the NR patient with progressive disease whereas tumor 227 is from the R patient with stable disease. The D-value for each tumor is averaged from those of PF TILs of this tumor. **(f).** ROC curve. The prediction accuracy of D value in distinguishing the tumors of non-responders from those of responders is AUC =100. 5-fold cross-validation supporting vector machine (SVM). AUC: The area under the ROC curve. ROC: Receiver Operating Characteristic. **(g).** Heatmap of ΔE values of the 8 exhaustion related markers by the TILs from D low to D high. 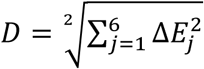, where ΔE value of each marker is the expression level difference between the PF TILs and their baseline non-front TILs. D value category is classified by D values of the PF TILs (**Online Methods**). Heatmap of another 3 exhaustion related markers is shown in **Extended Data Fig. 8b**.

Next, to explicit fundamentally different T cell functionalities as well as phenotypic gene signatures underlying D-high and D-low TILs, we analyzed another single-cell RNA-seq (scRNA-seq) dataset^28^ of 1,233 TILs pooled from 8 melanoma patients (**Online Methods**, **Extended Data Fig. 5**-**6**). Most of these patients received the ICBs of αPD-1, αCTLA-4, or αPD-1+αCTLA-4 (**Supplementary Table 4**). We used LAG3, TIGIT, HAVCR2 (TIM3), PDCD1 (PD-1), CTLA-4, IRF4, EOMES, and CD160, as exhaustion markers (**Fig. 2a**, **Extended Data Fig. 6**, **Supplementary Text**). We divided the PF TILs into four categories: D Extremely-high (top 1%), UQ_90 (D-high, top 2%-10%), UQ_75 (D-high, upper quartile excluded top 10%), and D-low (lower 75%), ref to **Fig. 2a**, **Supplementary Table 5**. D-high and D-low as well as non-PF TILs (TILs excluded by PFs) show distinct activation of T cell signaling pathways and functions, including cytotoxic T cell signaling, CD28 signaling, PCKe signaling, cytokine-related gene signature, and exhaustion gene signature (**Online Methods**, **Fig.2b-c**, **Extended Data Fig. 7**). It seems that non-PF TILs are resting T cells displaying neither exhaustion nor cytotoxicity^27^, but enriched with a central memory signature including markedly increased expression of IL-7R, LEF1, SELL, CCR7, and VAX2 (**Fig. 2d**). Most importantly, non-PF TILs have significantly low tumor-specificity described by CD39^low/neg^CD103^low/neg^^11,12^ (**Fig. 2e**). This explains why we observed a large portion of non-PF TILs that seem to be irrelevant to treatment response because they are tumor-infiltrating bystander T cells.

**Figure 2.**
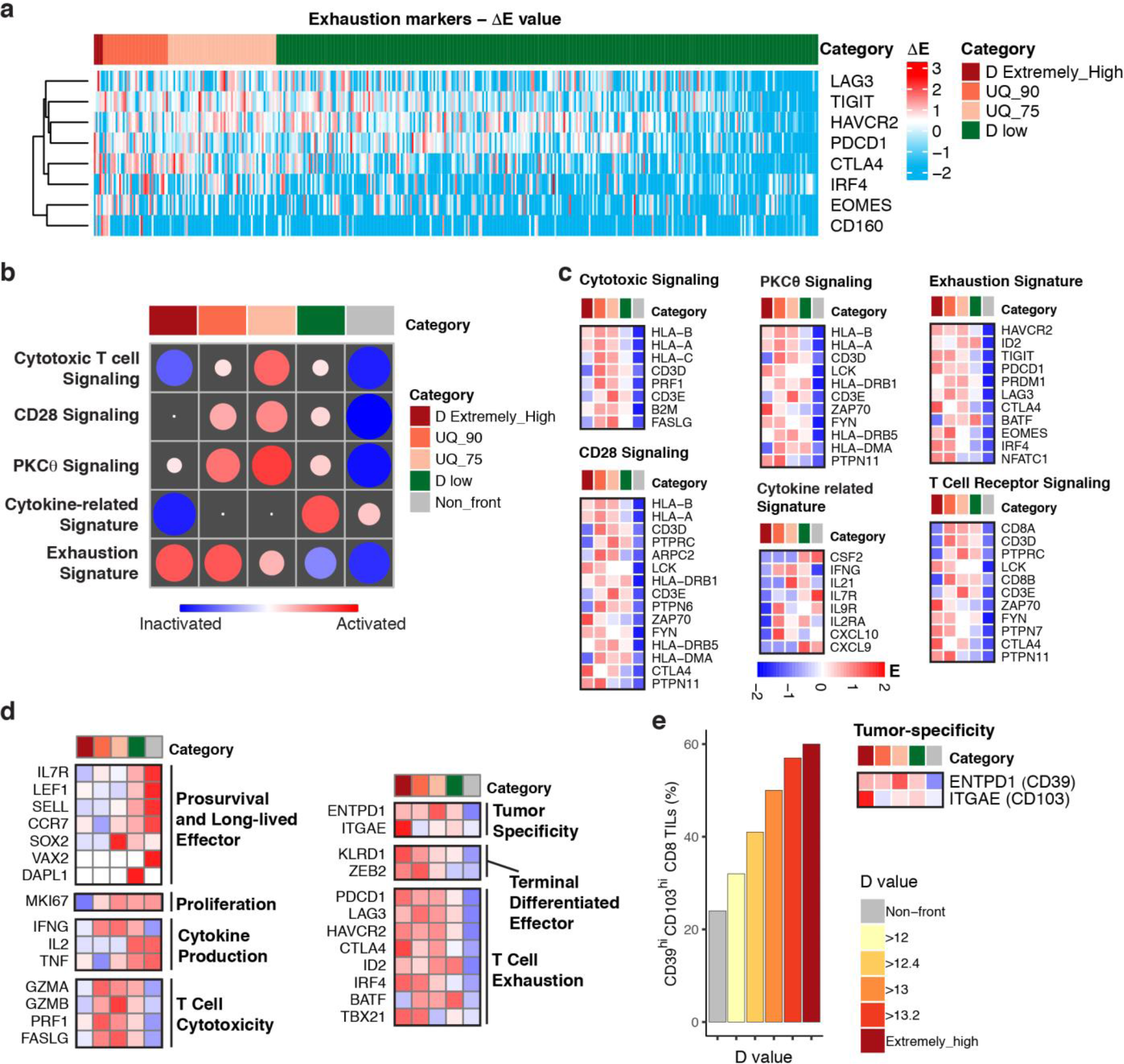
TILs with D low and D high show distinct activation of T cell signaling pathways and T cell functions. (a). TIL categories classified by D values from high to low. 401 PF TILs were classified into D Extremely-high (top 1%), UQ_90 (top 10% excluded D Extremely-high), UQ_75 (upper quartile excluded top 10%), and D low (lower 75%). The heatmap shows the ΔE values of the exhaustion markers of each PF TIL. D value of each PF TIL is derived from the ΔE values of the exhaustion markers. **(b).** Activated signaling pathways and signatures related to T cell functionality. Shown are the activation scores that evaluate the activation patterns of the T cell exhaustion related signaling pathways and signatures (**Online Methods**). **(c).** Heatmaps of expression levels of the signature genes associated with the activated signaling pathways. **(d).** Heat maps illustrating the relative expression of genes in different categories of TILs. Statistical significance is shown in **Extended Data Fig. 7**. **(e)**. Analysis of the tumor-specificity in TILs with different D-values. CD39 and CD103 are used to annotate tumor-specific TILs.

D-high and D-low TILs show fundamentally distinct signaling signatures. The TILs with D Extremely-high display highest exhaustion and inactivation of both cytotoxicity and effector functions, together with decreased IFN-g production. With the decrease of D values which comes with downregulation of T cell exhaustion, the tumor-specific TILs with UQ_90 and UQ_75 demonstrate a gradually upregulated activation of effector T cell signatures, including cytotoxicity and T cell activation through TCR, PCKe and CD28 signals. In addition, the less-exhausted D-low TILs are tumor-specific cytotoxic effectors (**Fig. 2b-c**). Of note, the TILs with D-high are terminally differentiated effector (KLRD1 and ZEB2), with the high expression of T cell exhaustion (PD-1, LAG3, CTLA-4, TIM3, etc.) and tumor specificity markers (CD39 and CD103), but inactivated in prosurvival and long-lived gene signature (e.g., IL7R, LEF1, and SELL), proliferation (Ki-67), cytokine production (INFG, IL2, and TNF), and cytotoxicity (GZMA, GZMB, PRF1, and FASLG). The result from scRNA-seq analysis is highly consistent with the observed D-high TILs in non-responders identified from the CyTOF data. The TILs with high D values possess their unique exhaustion accompanied by high tumor-specificity and irreversible T-cell dysfunction (at least not by αPD-1/αCTLA4), which lead to the therapy resistance of the present ICBs in non-responders.

To determine whether the good outcome of the clinical responders is determined by specific exhaustion markers after receiving αPD-1, αCTLA-4, or αPD-1+αCTLA-4, we further defined the contribution ratio of each marker to the D value, that is, DE_i_/E_i_, the ratio between increased expression level of each marker in a PF single cell compared to its non-PF baseline single cells (DE_i_) and the expression level of this marker in this PF single cell (E_i_), see **Online Methods**, **Extended Data Fig. 1**. Comparison of the contribution ratios of the exhaustion markers between responders and non-responders revealed that LAG3 has low contribution to D values in most PF TILs of responders (**Fig. 3a,c**). In other words, LAG3 does not show high expression levels in most PF TILs of responders. Analysis of the LAG3-low PF TILs by CD45RA, CD45RO, and CD127, revealed that these exhausted TILs show high effector but low memory function in the responders (**Extended Data Fig. 8a**). To predict efficacious ICB combinations based on the contribution ratios of the exhaustion markers, including PD-1, CTLA-4, LAG3, and TIM3, we defined an *in-silico* exhaustion value for each tumor by using the burst levels of DE/E and D values of the PF TILs (exhausted TILs), refer to **Online Methods**. The *in-silico* TIL exhaustion value, *Ex*, represents to what extent the selected exhaustion markers included in a new ICB combination have burst expression in D-high TILs after receiving αPD-1, αCTLA-4, or αPD-1+αCTLA-4 (**Online Methods**). By associating the predicted *Ex* values of tumors with their clinical responses to ICBs, we identified the important roles of LAG3 in developing alternative ICB strategies, e.g., αLAG3+αPD-1+αCTLA-4 (**Fig. 3b**). The role of LAG3 in ICB combinations remains to be identified^29,30^ though the combination of αLAG3 with αPD-1 or αCTLA-4 in ongoing clinical trials (NCT01968109).

**Figure 3.**
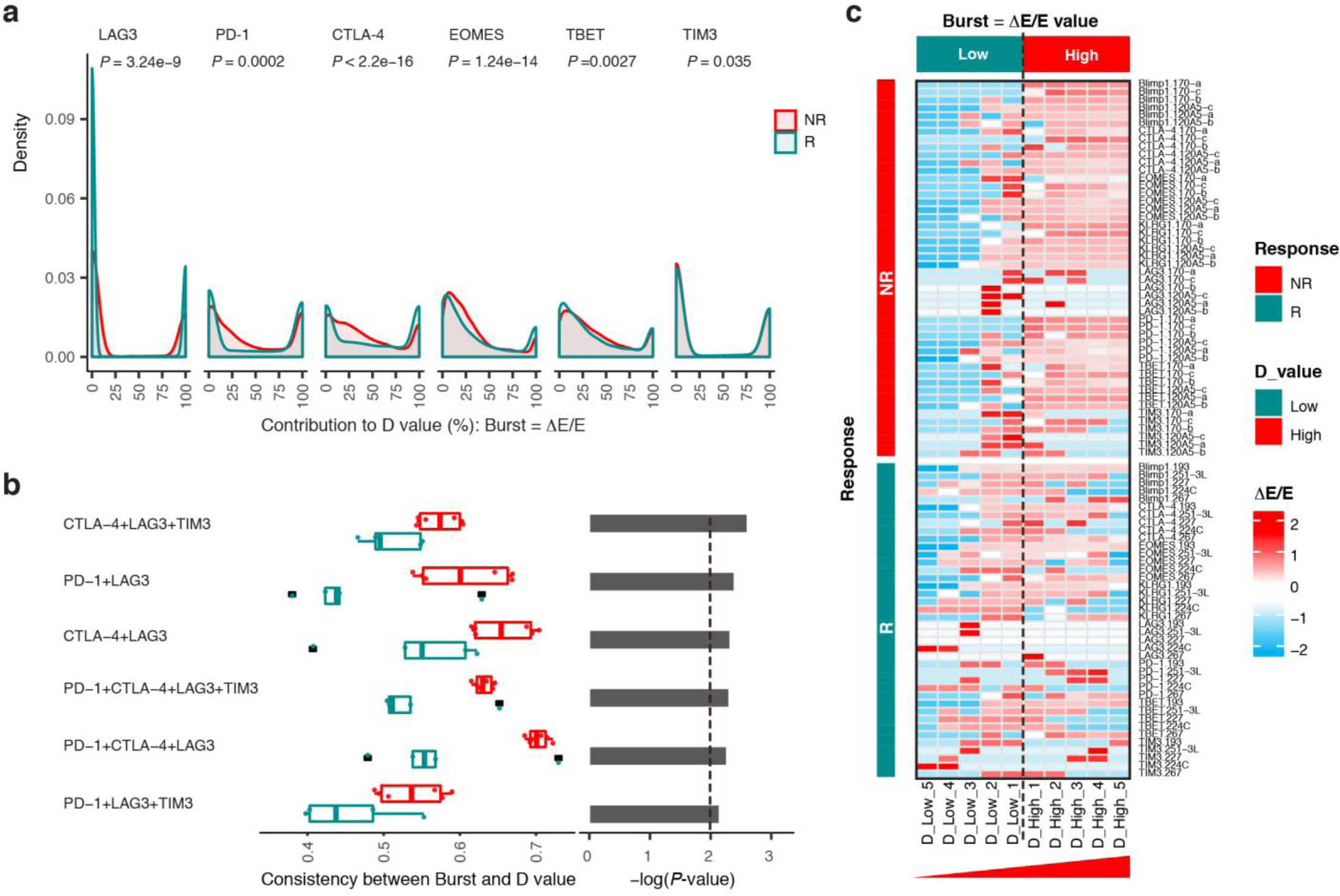
Burst expression of LAG3 and its important role in predicting alternative strategy for ICB. **(a).** Burst expression of the exhaustion markers included in human melanoma CyTOF data. Burst is defined as ΔE/E, describing to what extent the marker has burst expression change in the PF TIL after the ICB. ΔE is defined as in **Fig. 1f**, and E is the expression level of this marker in the PF TIL. 100 indicates that the burst is from 0 to a high level whereas 0 suggests no change. **(b).** Predicting efficacious immune checkpoint combinations. The candidate ICB combinations are determined by the consistency between the Burst values of the combined markers and D values of PF TILs. 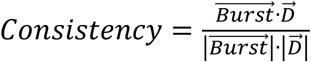. If the combined markers show significantly higher Bursts in D-high TILs of non-responders (P < 0.01, Mann-Whitney *U* test), the new alternative strategy takes the combined markers as a new ICB. The threshold is 2 for −log(*P*-value). **(c).** Heatmap of the Burst values of the 8 exhaustion related markers in non-responders (NR) and responders (R). Heatmap of another 3 exhaustion related markers is shown in **Extended Data Fig. 8c**.

Importantly, we validated the superiority of the new ICB combination, αLAG3+αPD-1+αCTLA-4, in eradicating established mice tumors and prolonging mice survival, compared to αPD-1+αCTLA-4 and other ICB combinations with αPD-1+αCTLA-4. By *in vitro* experiments, we firstly observed significantly upregulated expression levels of PD-1, CTLA-4, and LAG3 in the activated CD8^+^ T cells after co-culture with MC38-OVA tumor cells (**Fig. 4a**, *P* < 10^−16^, Two-sample Kolmogorov-Smirnov test), but no changes in the presence of αPD-1 + αCTLA-4 or isotype-matched antibody (IgG), see **Fig. 4b**. We next tested the antitumor capacity of αLAG3+αPD-1+αCTLA-4, compared to PBS, IgG, αLAG3, αPD-1+αCTLA-4, in the CT26 mouse model that shows resistance to most known ICBs. As expected, PBS, IgG, αLAG3, αPD-1+αCTLA-4 show resistance although moderate improvement for αPD-1+αCTLA-4 (**Fig. 4c**). Strikingly, αLAG3+αPD-1+αCTLA-4 eradicated the large established tumors with long-term tumor-free survival (**Fig. 4c-e**). Additionally, we also tested the antitumor ability of other ICB combinations, e.g., αTIM3+αPD-1+αCTLA-4 and αTIGIT+αPD-1+αCTLA-4, but no better improvement than αPD-1+αCTLA-4 treatment was observed (**Extended Data Fig. 10a**). Mechanistically, αLAG3+αPD-1+αCTLA-4 leads to markedly increased tumor-infiltrating CD4^+^ and CD8^+^ T cells, compared to other treatments (**Fig. 4f**). The statistical analysis by normalizing T cell number to per mg tumor demonstrates a dramatical increase of CD4^+^ and CD8^+^ T cells in the treatment by αLAG3+αPD-1+αCTLA-4 (**Fig. 4g**). Lastly, we implemented high-dimensional single-cell analysis by CyTOF on the tumors from the 5 treatment groups (**Online Methods**, **Extended Data Fig. 10b**). Notably, the T cell populations from αLAG3+αPD-1+αCTLA-4 show extremely high expression levels of GzmB (4 times higher than αPD-1+αCTLA-4 and ∼300 times higher than other treatments) and high tumor specificity, refer to **Fig. 4h** and **Extended Data Fig. 10b**.

**Figure 4.**
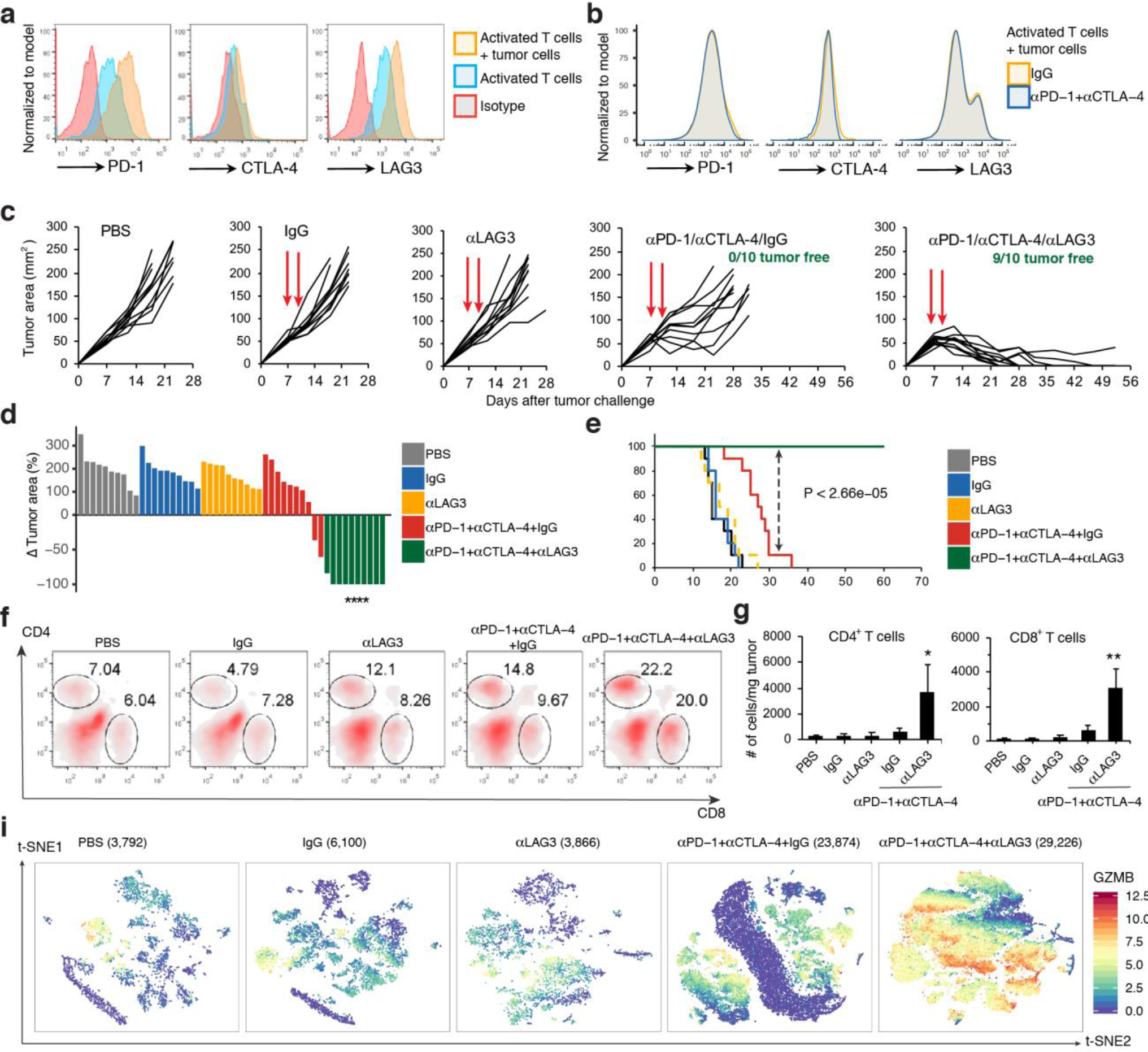
Efficacy of the new ICB strategy of αPD-1+αCTLA-4+αLAG3 in murine tumor models. **(a).** Flow cytometric analysis of αPD-1, αCTLA-4 and LAG3 expression on the activated CD8 T cells before and after cocultured with MC-38-OVA cells *in vitro* (n = 3). **(b).** Flow cytometric analysis of αPD-1, αCTLA-4 and LAG3 expression on activated T cells which cocultured with tumor cells in the presence of αPD-1 + αCTLA-4 or isotype-matched antibody (IgG) (n = 3). **(c-e).** CT26 tumor growth (c) in BALB/c mice that were treated with PBS or IgG or αLAG3 or αPD-1+αCTLA-4 or αPD-1+αCTLA-4+αLAG3, as indicated by arrow, and percent change in tumor volumes, **** *P* < 10^−9^, Fisher’s exact test (d) between days 7 and 49, and Kaplan Meier survival plot of different treatment groups (n = 10), log-rank test (e). **(f)**. Flow cytometric analysis of the percentage of CD4 and CD8 T cells in the tumor of mice treated with PBS or IgG or αLAG3 or αPD-1+αCTLA-4 or αPD-1+αCTLA-4+αLAG3 (n = 3). **(g)**. The cell numbers of CD4 and CD8 T cells in the tumors of mice treated with PBS or IgG or αLAG3 or αPD-1+αCTLA-4 or αPD-1+αCTLA-4+αLAG3 (n = 3). * P < 0.01, ** P < 0.001, Student’s *t* test. **(h)** t-SNE plot of CD45^+^ cells overlaid with the expression of GzmB by CyTOF data.

Overall, our high-dimensional single-cell based computational model provides a comprehensive evaluation of CD8^+^ TIL exhaustion by modeling CD8^+^ TILs in a high-dimensional expression space. The new computational biomarker, D value, plays a key role in predicting clinical response to ICB and discovering alternative efficacious ICB combination. Detailed investigation of the D-values by the marker contributions led to the observation of low expression of LAG3 in the clinical responders and discovery of the new combination of αLAG3 with αPD-1+αCTLA-4 that aims to reverse the LAG3-high TILs in non-responders. We validated the efficacy of the new ICB in a remarkable murine model for cancer immunotherapy. High-dimensional single-cell analysis by CyTOF displayed the dramatically increased GzmB in most TILs after αLAG3+αPD-1+αCTLA-4. Our results demonstrate the importance of characterizing and modeling of CD8^+^ TIL exhaustion by the high-dimensional markers representing fundamentally distinct types of T cell functions at the single-cell level.

## Methods

### Cell lines

CT26 and MC38 murine colon carcinoma cells were purchased from ATCC. MC38-OVA cells were generated by transducing MC38 cells by OVA-encoding lentivirus. Cells were cultured in RPMI 1640 Medium (Invitrogen) supplemented with 10% heat-inactivated fetal bovine serum (Thermo Scientific), 100 U/ml penicillin-streptomycin, and 2 mM L-glutamine (both from Invitrogen).

### Mice

BALB/c (Stock No: 000651), C57BL/6-Tg(TcraTcrb)1100Mjb/J (Stock No: 003831| OT-1) mice were purchased from The Jackson Laboratory. Male and female 6- to 8-week-old mice were used for each animal experiment. All experiments complied with protocols approved by the Institutional Animal Care and Use Committee at the Wake Forest School of Medicine.

### Reagent

IgG control, αPD-1 (clone RMP1-14), αCTLA-4 (clone UC10-4F10-11), αLAG3 (clone C9B7W), αLAG3 (clone RMT3-23), and αTIGIT (clone 1G9) were purchased from BioXcell. Human IL-2 was purchased from R&D Systems.

### *In vitro* cell coculture

Naïve CD8^+^ T cells were purified from the spleens of OT-I mice by isolation kit (STEMCELL Technologies, Cat#19858) according to the manufacturer’s protocol. OT-I-specific naïve CD8 T cells were cultured for 5 days with Dynabeads (cat#11452D, Thermo Fisher) in the presence of hIL-2(100 U/ml). After cultured for 5 days, the activated CD8 T cells were cocultured with 0.3×10^6^ MC38-OVA cells plus control IgG or PBS or MC38-OVA cells plus αPD-1 and αCTLA-4 for 48 hours in the presence of IL-2 (50 U/mL).

### Flow cytometry

FITC-, PE- or eFluor-conjugated mAbs (1:100 dilution) were used for staining after Fc blocking. Samples were acquired with Fortessa flow cytometer and data were analyzed with Flowjo software.

### *In vivo* murine tumor experiments

BALB/c mice were inoculated subcutaneously (s.c.) at the right flank with 1×10^6^ CT26 tumor cells in 0.1 mL of PBS. Treatment was started at day 7 when tumors area reached about 50 mm^2^ (∼8×7 mm). IgG control (100 µg, ∼4 mg/kg), αPD-1 (100 µg, 4 mg/kg), αCTLA-4 (100 µg, 4 mg/kg), αLAG3 (100 µg, 4 mg/kg), αTIM3 (100 µg, 4 mg/kg), and αTIGIT (100 µg, 4 mg/kg), alone or in combination, were intraperitoneal (i.p.) injection on days 7 and 10 after tumor injection. Tumors were measured by caliper and tumor area were calculated as width × length. At the time of sacrifice for analysis, mice were euthanized using CO_2_ and subsequent cervical dislocation.

### Mass cytometry antibodies

Metal conjugated antibodies were purchased from Fluidigm or conjugated to unlabeled antibodies in our lab. All non-platinum conjugations were performed using Maxpar Antibody Labeling Kit (Fluidigm) according to the manufacturer’s protocol and were performed at 100 mg scale. Antibodies include: 89^Y^-CD45, 141^Pr^-Gr1, 142^Nd^-CD39, 145^Nd^-CD69, 145^Nd^-CD8a, 147^Sm^-CD160, 148^Nd^-Ki67, 149^Sm^-CD19, 150^Nd^-CD44, 151^Eu^-CD25, 152^Sm^-CD3e, 153^Eu^-PD-L1, 154^Sm^-CTLA4, 155^Gd^-TIGIT, 156^Gd^-KLRG1, 158^Gd^-Foxp3, 159^Tb^-2B4, 160^Gd^-Tbet, 162^Dy^-TIM3, 164^Dy^-CD62L, 165^Ho^-IFNg, 167^Er^-Nkp46, 169^Tm^-VISTA, 170^Er^-EOMES, 171^Yb^-GzmB, 172^Yb^-CD11b, 174^Yb^-LAG3, 175^Lu^-CD127, 176^Yb^-ICOS, 209^Bi^-CD11c and 194^Pt^-Live/Dead. Every antibody was used at 1:100 dilution on samples.

### Mass cytometry analysis of murine tumors

Tumors were dissected, manually dissociated, and digested enzymatically with Collagenase D (Sigma) and DNase I (Roche) in PBS containing 2% FBS for 20 min at room temperature with continuous agitation. Add EDTA to a final concentration of 10mM and incubate at room temperature for 5 min. Pour the entire suspension through a 70 um filters into RPMI-1640 supplemented with 10% FBS. Filtered cells were then washed twice with FACS buffer and total cell concentration determined using an automated cell counter (Thermo Fisher). 2 × 10^6^or fewer cells per tumor were performed by cell surface staining, cytoplasmic antigen staining, and nuclear antigen staining according to the manufacturer’s protocol (Fluidigm). Samples were shipped to the CyTOF Core of Dana-Farber Cancer Institute and then analyzed using a CyTOF2.

## Methods

### CyTOF and scRNA-seq datasets

We used three single-cell datasets from CyTOF and scRNA-seq platforms in our analyses, that is, human melanoma CyTOF data, mouse melanoma CyTOF data, and human melanoma scRNA-^5^seq data. Human melanoma CyTOF data include 14 specimens from 7 patients (n =11, tumor tissues) and 3 healthy donors (n =3, PBMC). These patients received αPD-1, αCTLA-4, or αPD-^7^1+ αCTLA-4, **Supplementary Table 1**. The responses to ICB are progressive disease or PD, partial response or PR, and stable disease or SD. The PBMC samples from the 3 healthy donors are the baseline control for the ICB. The specimens for the responses to ICB are PD (n =6 from 2 patients), PR (n =1 from 1 patient), and SD (n =4 from 4 patients). Mouse melanoma CyTOF data include 52 tissue specimens (n =52) from 34 mice received αPD-1+GVAX or αCTLA-4+GVAX (n =34) and 18 control mice that only received GVAX tumor vaccine (n =18), **Supplementary Table 2**. The tumor volumes of the 34 mice after ICB therapy and the 18 control mice were derived. Human melanoma scRNA-seq data include 19 tumor tissue specimens from 19 patients, in which 8 patients (n =8) have more than 50^CD8^+ TILs. These patients received αPD-1, αCTLA-4, or αPD-1+ αCTLA-4, **Supplementary Table 4**.

The CyTOF data were bead-normalized and debarcoded mass cytometry. We applied Flow-17 SOM analysis to the raw data. The expression values were arcsinh transformed by the antibodies for transformation (**Supplementary Tables 5-6**). In order to apply HD-scMed to the tumors and compare the tumors, we further applied upper-quartile normalization to the processed CyTOF data. The CD8^+^ TILs were identified by consensus clustering of the antibodies for cell linage (**Supplementary Tables 5-6**, **Extended Data Figs. 2-3**). The scRNA-seq data were the processed data that were normalized by house-keeping genes as described in ref.

### High-dimensional single-cell based computational method (HD-scMed)

HD-scMed identifies exhausted TILs from a high-dimensional expression space defined by a variety of exhaustion markers, including immune checkpoints and related transcriptional factors (**Extended Data Fig. 1**). The number of exhaustion markers included in HD-scMed is determined by the available antibodies in the CyTOF data and the functionality of diverse exhaustion markers in the scRNA-seq data. The mathematical model utilized by HD-scMed is a well-established multiple-objective optimization model, Perato Optimization (PO). The key role of the PO model is to facilitate identification of a subset of^CD8^+ TILs with exhaustion in terms of all considered exhaustion markers within the high-dimensional expression space.

### Multiple-objective optimization model, Perato Optimization

A general multiple-objective optimization model (MOO) comprises a set of *n* decision variables, x, a set of *k* objective functions, y, and a set of constraints, e(x) ≤ 0. Objective functions and constraints are functions of the decision variables^31^.

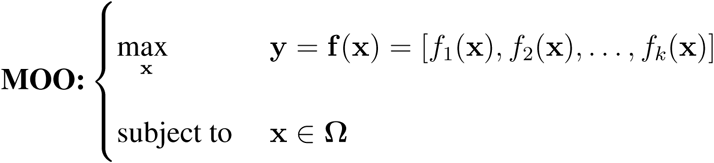

where Ω ={x =(x_1_, x_2_,…,x_*n*_) ∈ X|e(x) =[*c*_1_(x), *c*_2_(x), …,*c_m_*(x)] ≤ 0} is called as a feasible set, containing the set of decision vectors x that satisfies the constraints e(x) ≤ 0.

In this MOO model, decision variables, x, represent the CD8^+^ TILs that are coded by the molecules, genes or proteins, from Ω, objective values, y denote the expression levels of the exhaustion markers from these CD8^+^ TILs (x), available in the CyTOF or scRNA-seq data. But the issue is that the mathematical expression of the objective functions, f(x), are unknown. Alternatively, Pareto Optimization (PO) was considered.

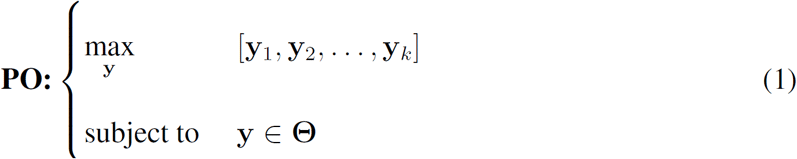

The PO model does not require the exact expression of the mathematical equation, f(x), in the MOO model. The variables, y, denote the expression levels of exhaustion markers, E, across the TILs. The feasible region of the PO model mapped from decision space X to the objective space Y, is denoted by Θ. The objective space, Θ, i.e., the high-dimensional expression space, contains the points defined by all candidate TILs of interest, each of which, 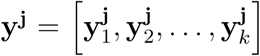, j ∈ X, represents the expression levels of the markers from a TIL j.

For the application of HD-scMed to human CyTOF data, the exhaustion markers (*i* =1, 2, …,6) are the ICs, PD-1, CTLA-4, LAG3, and TIM3, and the TFs, EOMES and TBET. In the PO model, each y^j^ is defined by the expression levels of the 6 exhaustion markers from a TIL j, j ∈ X. The expression levels of the TIL j, y_*j*_, is expressed as 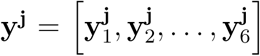, j ∈ X. For mouse CyTOF data, each y^j^ is defined by the exhaustion markers, αPD-1, TIM3, TIGIT, VISTA, EOMEs, and TBET. For human scRNA-seq data, we considered 15 well-studied exhaustion markers initially. But due to the diversity of the 15 markers in determining functionality of exhaustion, we identified 8 markers from 3 clusters of the 15 markers in the final model (**Supplementary Text**, **Extended Data Fig. 5**), that is, PDCD1 (PD-1), CTLA-4, LAG3, TIGIT, HAVCR2 (TIM3), IRF4, CD160, and EOMES.

### Pareto dominance

The PO model relies on a Pareto dominance relationship, ≻, defined for any two single cells in the decision space, Ω, to identify which TILs are exhausted. If two TILs have a ≻ b, or say, TIL a dominates TIL b, the expression levels of the *k* markers in a are higher than or equal to those of b, but a ≠ b. More precisely, we have

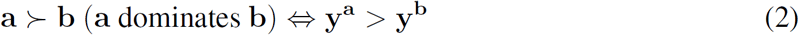

where y^a^ and y^b^ are the marker expression levels of the candidate TILs, a and b. If we have y^a^ > y^b^, we need the follows.

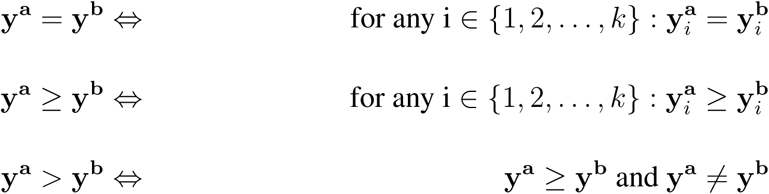

### Perato front or PF

In this paper, Perato front or PF is used to identify exhausted TILs in the high-dimensional expression space defined by the exhaustion markers. The PF TILs are also called nondominated TILs. Precisely, for a subset Ω′ ⊆ Ω, the function ℘(Ω′) defines the set of nondominated decisions in 4 Ω′:

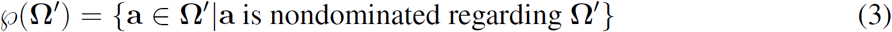

where

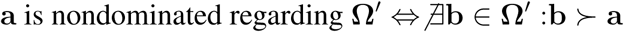

The ℘(Ω′) is the nondominated set with regard to Ω′. The set of mapped objective vectors, f(℘(Ω′)), is the Pareto-optimal front for Ω′. Moreover, the set X_℘_ =℘(Ω) is called the **Pareto-optimal set** and the set Y_℘_ =f(X_℘_) comprises the **Pareto-optimal** front, or for short, **Pareto front (PF)**. The PF contains the exhausted TILs in context of the *k*-dimensional exhaustion markers. The PF is a set of the expression level vectors, in which each vector denotes the expression levels of the markers from one single cell within the PF. In this paper, to simplify the terms, we called the exhausted TILs in the PF as PF TILs.

### Strength score and fitness score

Identification of the PF TILs, as shown in equation of (3), by HD-scMed requires strength score and fitness score defined by Pareto dominance. The detailed description of the algorithms used in PO model, such as Strength Pareto Evolutionary Algorithm 2 (SPEA2), can be found in Supplementary Text. Of note, our PO problem for identification of exhausted TILs is different from traditional PO problems defined by continuous variables or known objective functions, f(x). In our PO model, the decision space of the candidate TILs has a fixed number of *n*, which is the CD8^+^ TIL number in the CyTOF data or scRNA-seq data. However, the decision space Ω in the traditional PO problem was defined by continuous variables or known objective functions. This reality determines that the traditional PO problems cannot enumerate all decisions from decision space, requiring evolutionary algorithms, e.g., SPEA2.

Strength score and fitness score were designed by SPEA2 to evaluate which decisions should be included into PF (**Supplementary Text**). In our HD-scMed, evolutionary algorithms are not required but the strength score and fitness score are still needed to identify PF TILs. Based on SPEA2, we defined the strength score *S* and fitness score *F*.

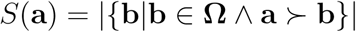

and

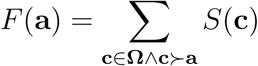

*S* and *F* scores make it feasible to quantify the exhaustion of TILs by the high-dimensional exhaustion markers. High strength score and low fitness score define exhausted TILs in this study. Thus, the PF TILs were defined as those TILs with strength score higher than zero and fitness score that equals to 0, i.e.,

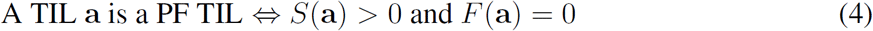

### D value

D value is designed for further categories of PF TILs, exploring the extents of exhaustion determined by the high-dimensional exhaustion markers. The D value for a PF TIL a is defined as the Euclidean distance or straight-line distance between a and its dominating non-front TILs, Ω″, where Ω″ satisfying that

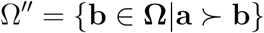

Precisely, D value for the TIL a is called *D*_a_,

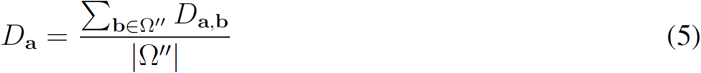

where *D*_a,b_ is defined by the ΔE_*i*_ of the exhaustion markers in the high-dimensional expression space, Ω. 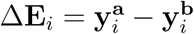.

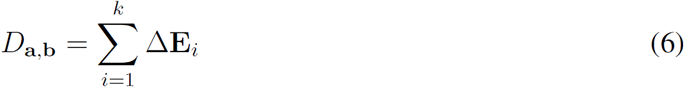

In the analysis of human melanoma CyTOF data (**Fig. 1**), the PF TILs were classified into D high and D low categories. *D* =1.25 is the threshold for D high and D low categories. *D* =0.25, 0.5, 0.75, and 1, are the additional thresholds for D low 5, D low 4, D low 3, D low 2, and D low 1. *D* =1.5, 2, 2.5, and 3, are the additional thresholds for D high 1, D high 2, D high 3, D high 4, and D high 5.

D value for a tumor, *t*, is defined by the D values of its PF TILs, 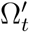. D value at the tumor 113 level, *D_t_*, is critical for predicting response to ICB.

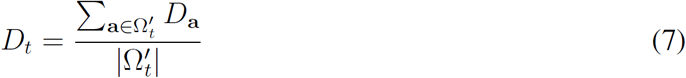

### Burst value

Burst value is extended from the ΔE value required for D value. The goal of Burst value is to investigate how the exhaustion is regulated by specific exhaustion markers after the tumor or the patient received the ICB therapy. The rationale behind Burst value is that resistance to ICB should be caused by the irreversible high exhaustion (at least not by αPD-1 or αCTLA-4) through regulating the burst expression of specific exhaustion markers. For a PF TIL, a, the Burst value for the marker *i*, is defined as *B*_a,*i*_,

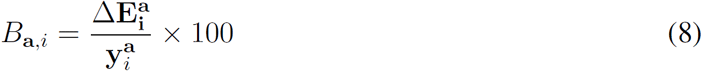

where 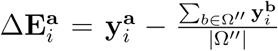, Ω″ ={b ∈ Ω| a ≻ b}. The Burst value ranges from 0 to 100. 0 where The Burst value ranges from 0 to 100. 0 means no expression change by comparing the PF TIL with its baseline non-front TILs whereas 100 indicates there is a burst from 0 to a high value for this exhaustion marker in the TIL after the tumor or the patient received the ICB therapy.

### Predicting responses to ICB by D values at the tumor level

As shown in **Fig. 1** and **Extended Data Fig. 4**, the D values at the tumor levels, *D_t_*, are highly associated or correlated with responses to ICB. This is because that D value highly represents the exhaustion of CD8^+^ TILs by considering the high-dimensional exhaustion markers. To further evaluate the importance of D values in predicting responses to ICB, we constructed the prediction model powered by a 5-fold cross-validation supporting vector machine (SVM). The *x* variables are the D values from the tumors, and the *y* variables for predictions are the responses to ICB from human patients or the tumor volumes derived from the mice after receiving ICB. The prediction performance is evaluated by the Receiver operating characteristic (ROC) curves. The prediction accuracy is described as the Area Under the Curve (AUC), which ranges from 50% to 100%.

### Predicting new combination strategy for ICB by both D values and Burst values of PF TILs

To figure out which exhaustion markers have burst expression in the PF TILs after the tumor received ICB and how these markers contribute to the responses to ICB, we developed a computational strategy to evaluate the consistency between the D values and Burst values of a combination, C, of specific exhaustion markers, that is, the *in silicon* exhaustion value, *Ex*. For a tumor, *t*, and its PF TILs, 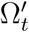, the consistency between D values and Burst values for the PF TILs, 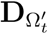 and 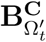, is defined as follows.

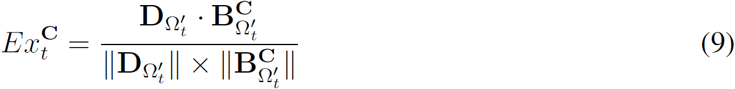

where 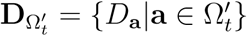 and 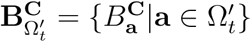.

With the changes of the combinations of exhaustion markers, *i* =1, 2, …, *k*, the *B*_a_ values for the marker combinations are updated. For a marker combination, C, the *B*_a_ is defined as follows.

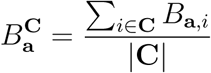

As shown in **Fig. 3b**, the prediction of the alternative strategy for ICB combination is based on the association of the exhaustion values, 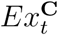, and the responses to ICB in both responders and non-responders. This prediction was implemented to human melanoma CyTOF data only. Precisely, the exhaustion values are 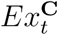 (*t* ∈ NR and R), of a marker combination, C, where NR includes the tumors from PD response and R is the set of the tumors from PR and SD responses. The significant exhaustion marker combinations were predicted by the statistical *P* value from the association between the exhaustion values and ICB responses. *P* < 0.01, Mann−Whitney *U* test.

### Pathway or gene signature activation

The pathway analyses were implemented by the Ingenuity Pathway Analysis (IPA) software (https://www.qiagenbioinformatics.com/products/ingenuity-pathway-analysis/). The statistical *P* values identified for the enriched signaling pathways are the p-values from Fisher′s exact test that are adjusted by Benjamini-Hochberg (B−H) method. The activation scores for pathways and gene signatures were calculated as follows,

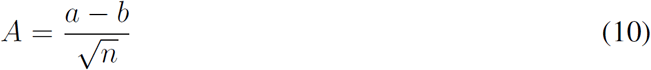

where *a* is the number of upregulated genes,*b* is the number of downregulated genes, and *n* is the total gene number in the pathway or gene signature.

### Statistical analyses

We have implemented the statistical analyses by R. Mann−Whitney *U* test calculates the two-sides p-values. Pearson product-moment correlation coefficient test returns the 2-tailed p-values. Analysis of Variance (ANOVA) test is carried out by one-way ANOVA. The *P* values are from the *F*-distribution. A multiple-testing corrected *P* value,*q* value, is calculated using the Benjamini-Hochberg method for the DEGs calculated by Student′s *t* test. The DEGs were defined by *P* value less than 0.001 and corrected p-values less than 0.01. The corrected p-values by Benjamini-Hochberg (B−H) method were also used for the Fisher′s exact test employed for the pathway analysis in IPA software. The enriched signaling pathways were identified by B−H *P* values with a threshold of 0.05. The *P* values from Fisher′s Exact Test is right-tailed in the IPA software.

### Software availability

Software used to generate all analyses in this manuscript is publicly available as a Python package, HD-scMed (https://github.com/guangxujin/HDscMed) and included here as Supplementary Software. HD-scMed can be accelerated by GPU computing or high-performance computing (HPC).

### Data availability

Full CyTOF data sets, scRNA-seq data set, and command lists to identify the POFs and the quantitative values are included as **Supplementary Data 1-3**. The published data used in this study can be accessed in the Gene Expression Omnibus under accession numbers GSE72056, and the Flow repository (https://flowrepository.org/) under accession FR-FCM-ZY6C and FR-FCM-ZY6A.

## Acknowledgements

Put acknowledgements here.

## Author contributions

**Extended Data Fig. 1.**
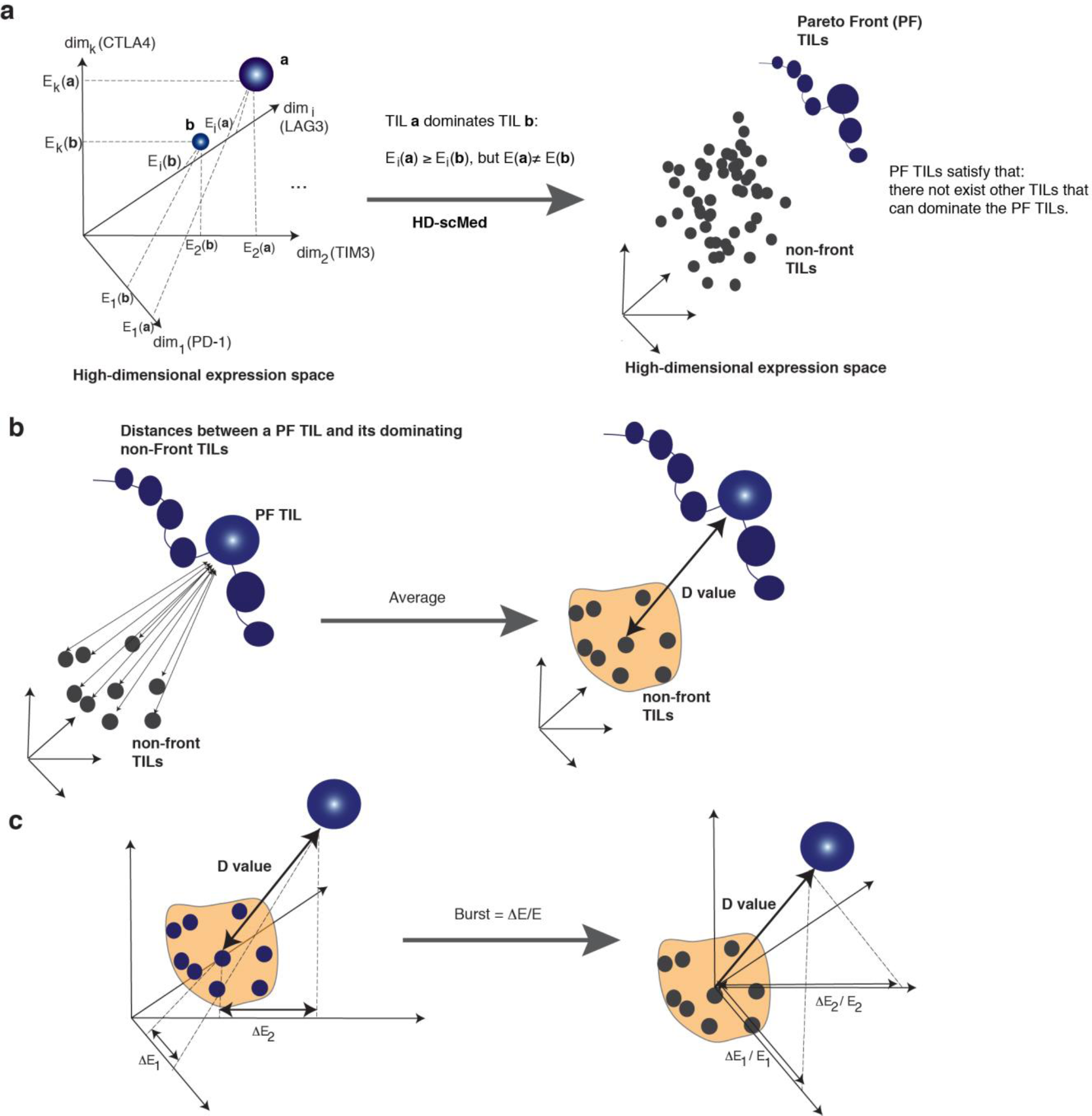
High-dimensional single-cell-based computational method (HD-scMed) for evaluation of TIL exhaustion within a high-dimensional expression space. **(a).** The high-dimensional expression space comprises the expression levels of TILs by a variety of exhaustion markers (PD-1, αCTLA-4, LAG3, TIM3, etc.) with their related transcriptional factors (e.g., EOMES and TBET). To quantify which TILs are exhausted in terms of the expression levels of the high-dimensional markers, the exhausted TILs are evaluated by Pareto dominance defined by a multiple-objective optimization (MOO) model (**Online Methods, Supplementary Text**). Pareto dominance from the MOO model identified a subset of TILs, called as Pareto Front (PF) TILs, each of which are optimal after trading-off the expression levels of high-dimensional exhaustion markers (by Strength score and Fitness score as described in **Online Methods**). A PF TIL is optimal because no other TILs can dominate it (**right panel**). Other than PF TILs, the TILs dominated by PF TILs are non-front TILs, the baseline for the exhaustion level of the PF TILs. **(b).** D value is calculated for each PF TIL and its dominated non-front TILs. Euclidean distance or straight-line distance between each PF TIL and each of its dominated non-front TIL is calculated. D value at the tumor level is defined as the average value of the Euclidean distances between the PF TIL and its dominated non-front TILs. D value is an integrative quantification criterion for TIL exhaustion by considering the high-dimensional expression information of the exhaustion markers. **(c).** Two critical values, ΔE and ΔE/E (Burst), were derived from D value for each exhaustion marker. The relationship between ΔE and D value is that 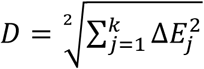, where k is the number of dimensions or the exhaustion markers (**Online Methods**). The Burst value is essential for evaluating which markers have burst expression (from a low level to a high level) after receiving the ICB, e.g., αPD-1, αCTLA-4, and αPD-1+αCTLA-4.

**Extended Data Fig. 2.**
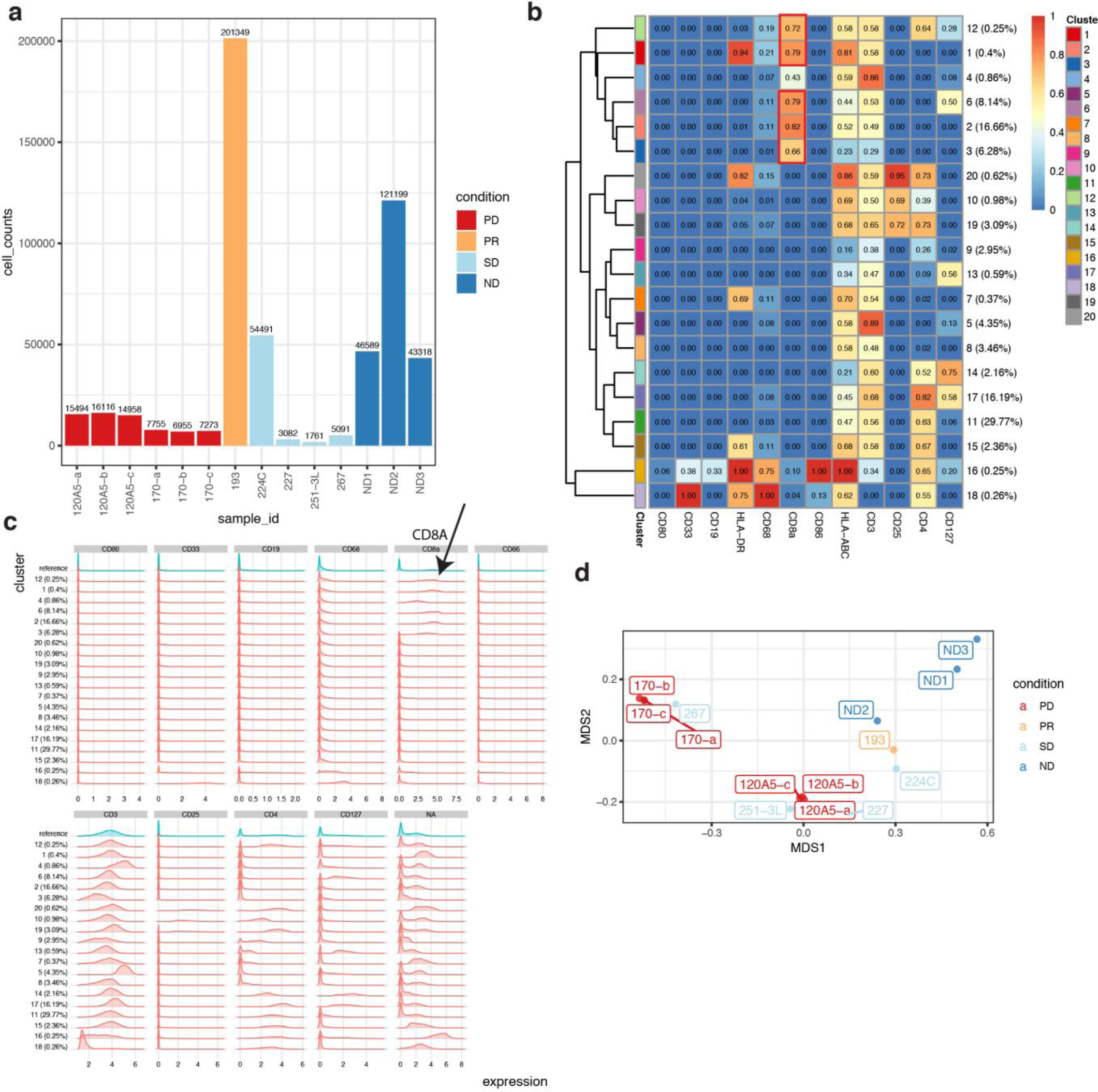
Identification of CD8^+^ TILs from human melanoma CyTOF data. **(a).** Cell counts for the specimens. Sample annotation is same as **Fig. 1**. **(b).** Consensus clustering of the pooled TILs from the 14 specimens, following FlowSOM analysis pipeline. The cluster # is set as 20. The CD8^+^ TILs are identified as those clusters with normalized CD8A expression level higher than 0.50 (**Online Methods**). **(c).** Histogram of clustering distribution. X axis is the expression level of the markers and y axis is the distribution density along the clusters. **(d).** Evaluation of the batch effect of the CyTOF data by multidimensional scale (MDS). Batch effect removal for the analyses in (**a-c**) is included in **Online Methods**.

**Extended Data Figure 3.**
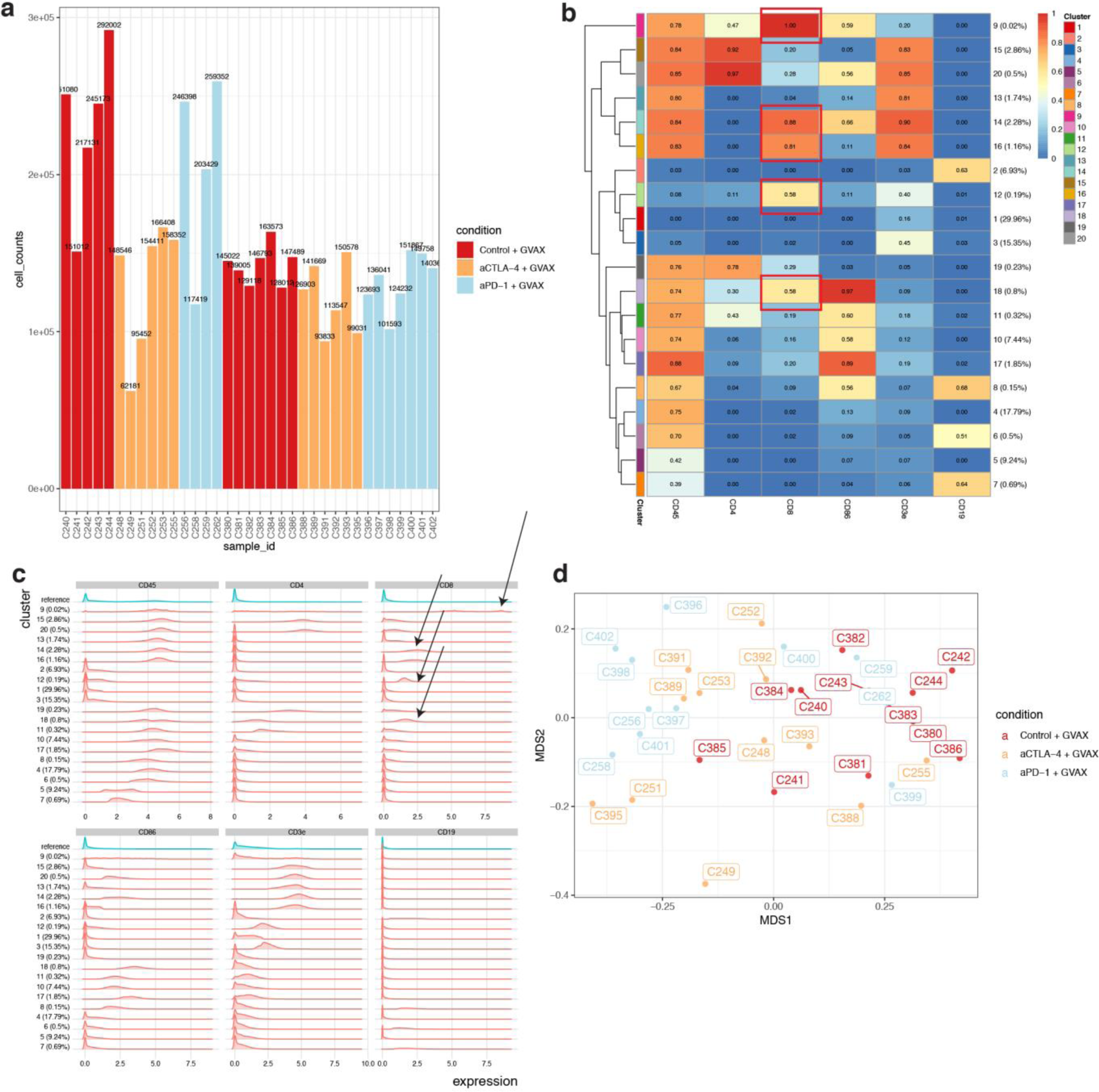
Identification of CD8^+^ TILs from mouse melanoma CyTOF data. **(a).** Cell counts for the specimens. Sample annotation is Shown in **Supplementary Table 2**. **(b).** Consensus clustering of the pooled TILs from the 52 specimens, following FlowSOM analysis pipeline. The cluster # is set as 20. The CD8^+^ TILs are identified as those clusters with normalized CD8 expression level higher than 0.50 (**Online Methods**). **(c).** Histogram of clustering distribution. X axis is the expression level of the markers and y axis is the distribution density along the clusters. **(d).** Evaluation of the batch effect of the CyTOF data by multidimensional scale (MDS). Batch effect removal for the analyses in (**a-c)** is included in **Online Methods**.

**Extended Data Figure 4.**
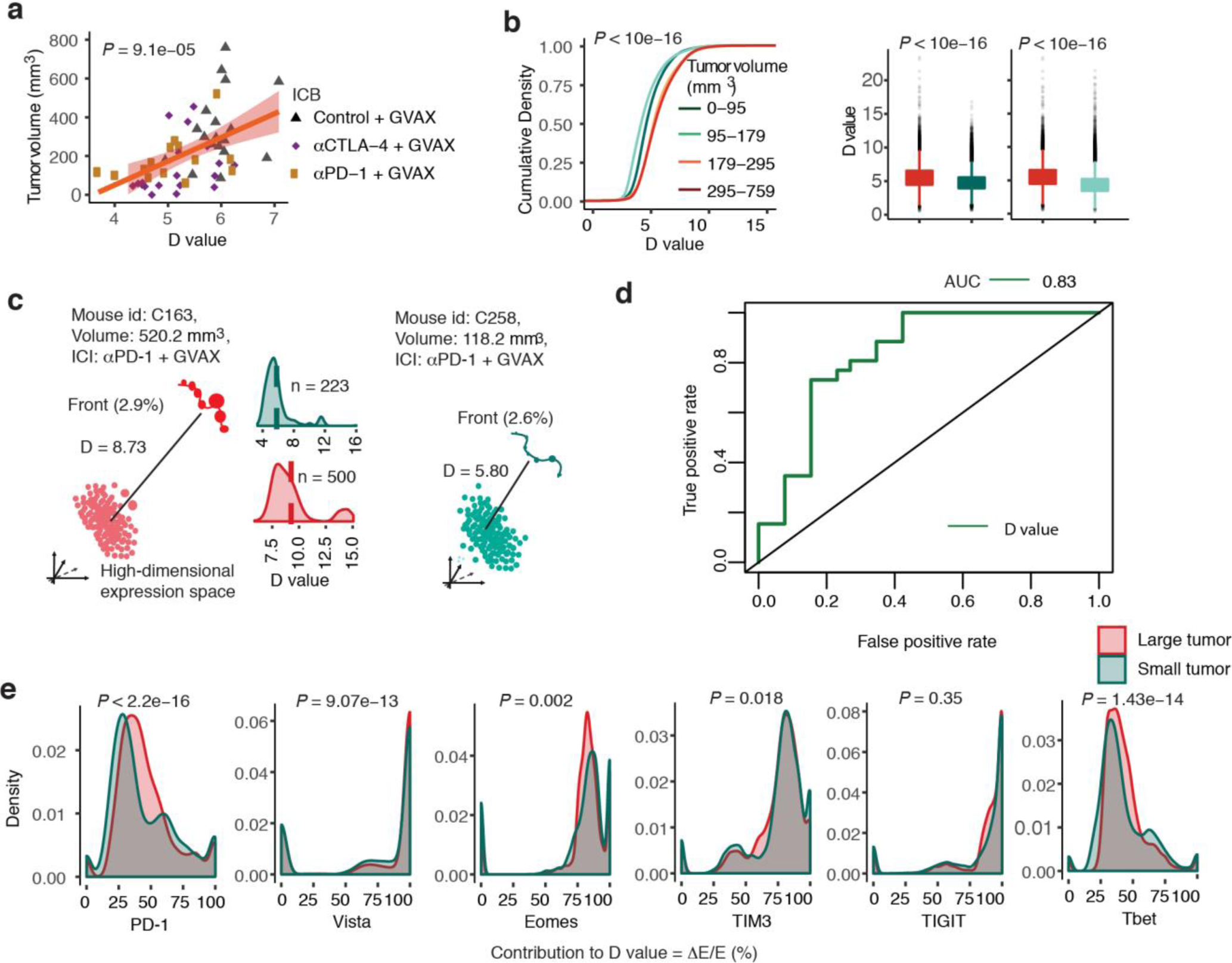
D value from HD-scMed in distinguishing the responses to ICB in 34 melanoma mice. **(a).** Correlation between D values derived from the PF TILs of the tumors and the tumor volumes after receiving the ICB therapies. **(b)**. The cumulative distributions of the D values in the PF TILs from the mouse group with different tumor volumes. P < 10e−16, Mann-Whitney *U* test. **(c).** An instance to show the difference of D values in two mice with different sizes of tumors after receiving αPD-1 + GVAX. The first mouse, C163, has a large tumor (within the first quartile of tumor sizes) whereas the second mouse, C258, bears a small tumor within the fourth quartile. **(d)**. ROC curve. The prediction accuracy of D value in distinguishing the tumors of non-responders from those of responders is AUC = 0.83. 5-fold cross-validation supporting vector machine (SVM). **(e).** The comparison of burst values of the 6 exhaustion markers available in the melanoma mouse CyTOF data.

## Supplementary Text

**Extended Data Figure 5.**
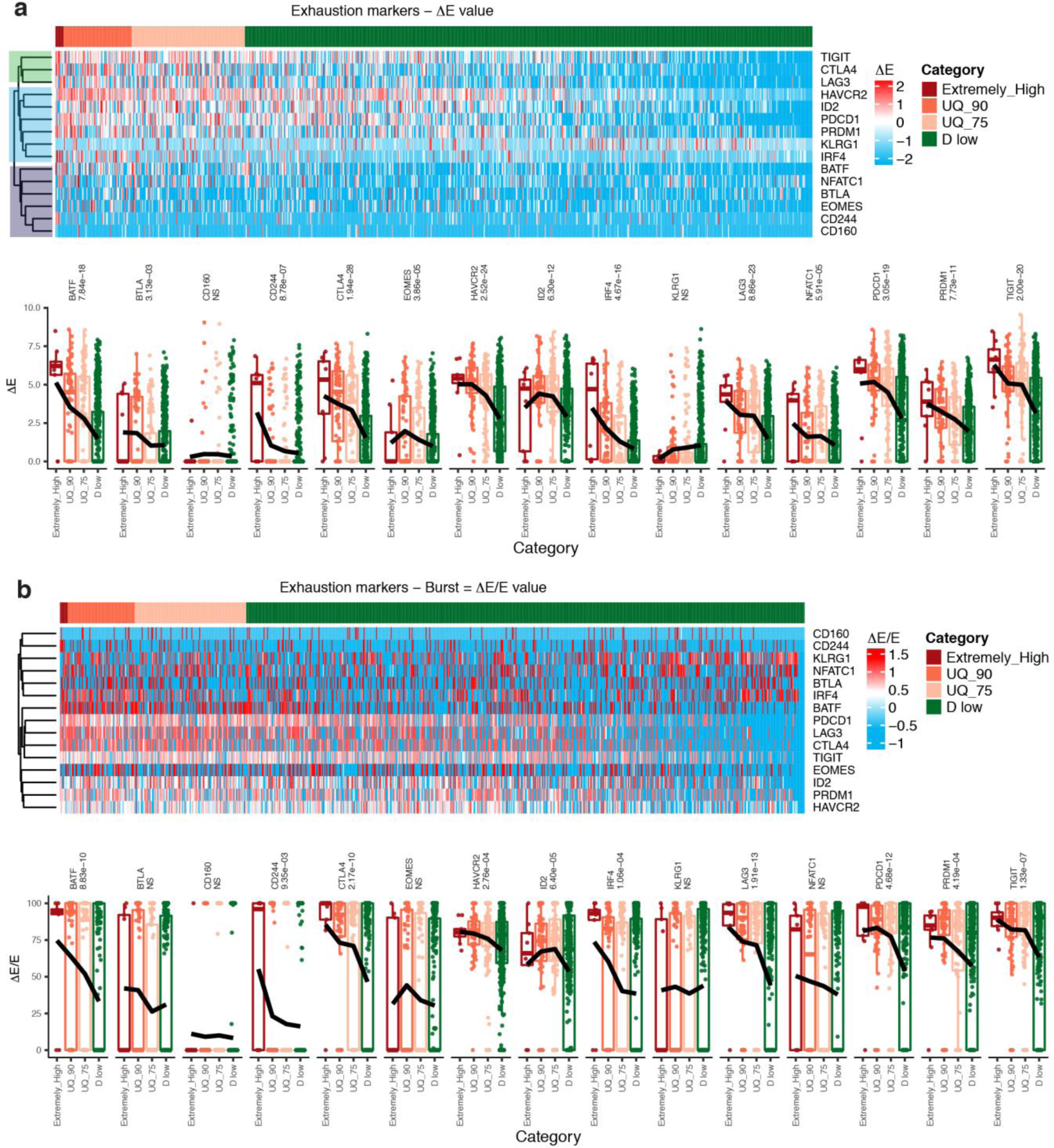
Hierarchical clustering of 15 exhaustion markers by ΔE and Burst values. **(a).** Hierarchical clustering of 15 exhaustion markers by ΔE values (top panel). The PF TILs were classified into four categories as defined by the percentiles in **Fig. 2**. Lower panel shows the statistical significance of the difference of ΔE values among the 4 TIL categories. One-way ANOVA analysis. (b) Hierarchical clustering of 15 exhaustion markers by Burst values (top panel). Lower panel shows the statistical significance of the difference of ΔE/E values among the 4 TIL categories. One-way ANOVA analysis.

**Supplementary Text:** The hierarchical clustering of the 15 exhaustion markers suggests their diverse roles in determining TIL exhaustion functions (**Extended Data Fig. 5**). The 15 markers were divided into 3 clusters. As an example, KLRG1 shows distinct pattern of ΔE values, compared to other markers. We also clustered the 15 exhaustion markers by their burst values. By combined the two clustering results, we selected 8 markers by satisfying (1) covering all 3 clusters of **Extended Data Fig.5a**; (2) covering the top significant markers shown in the clusters; (3) the NS markers showing the same trend as D value (CD160).

**Extended Data Figure 6.**
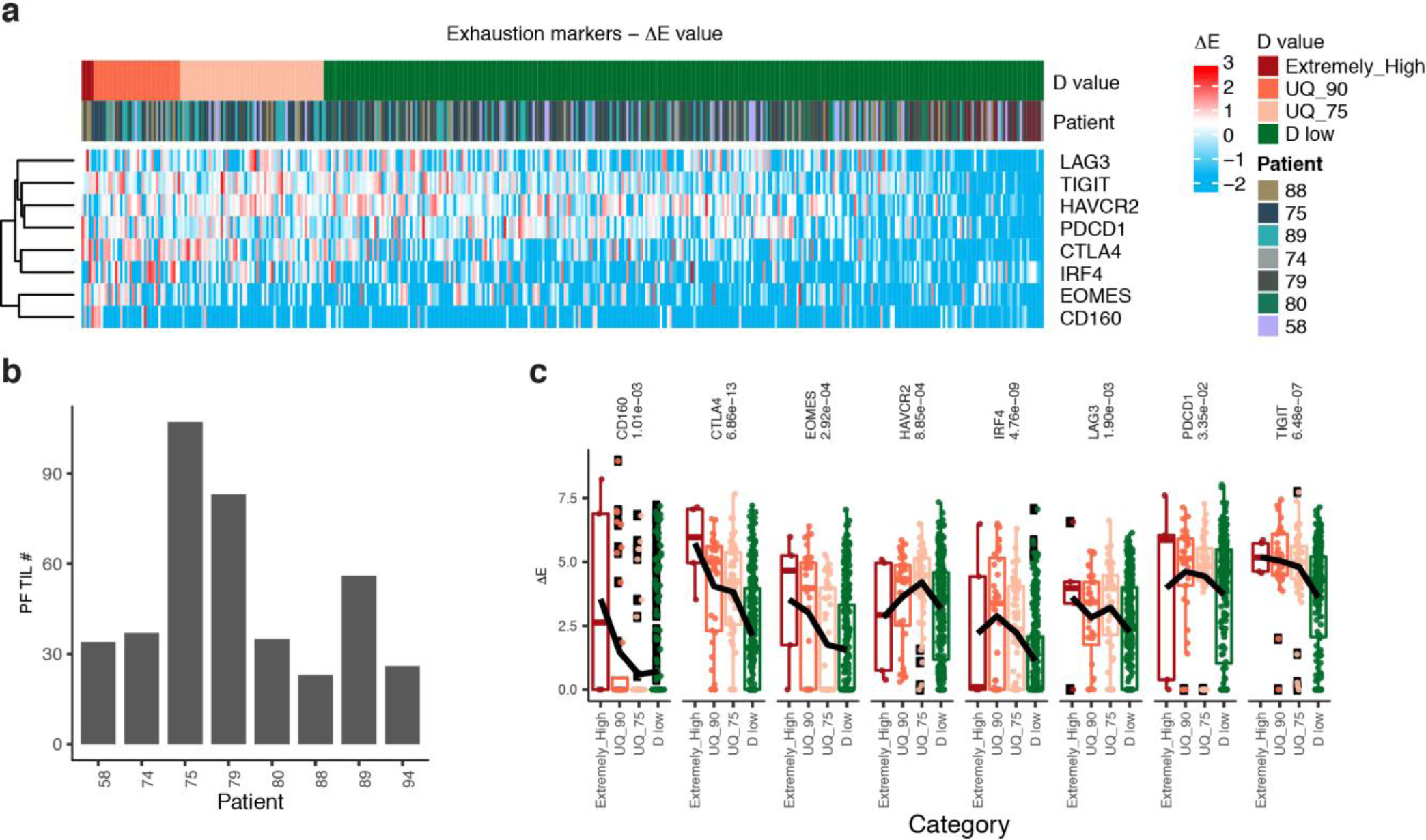
**(a).** Patient information added as column annotation. **(b).** the numbers of PF TILs in melanoma patients. **(c).** The statistical significance of the difference in ΔE values of these 8 markers among the identified four TIL categories. One-way ANOVA analysis. Patient information with PF TIL numbers in the Fig.2a.

**Extended Data Figure 7.**
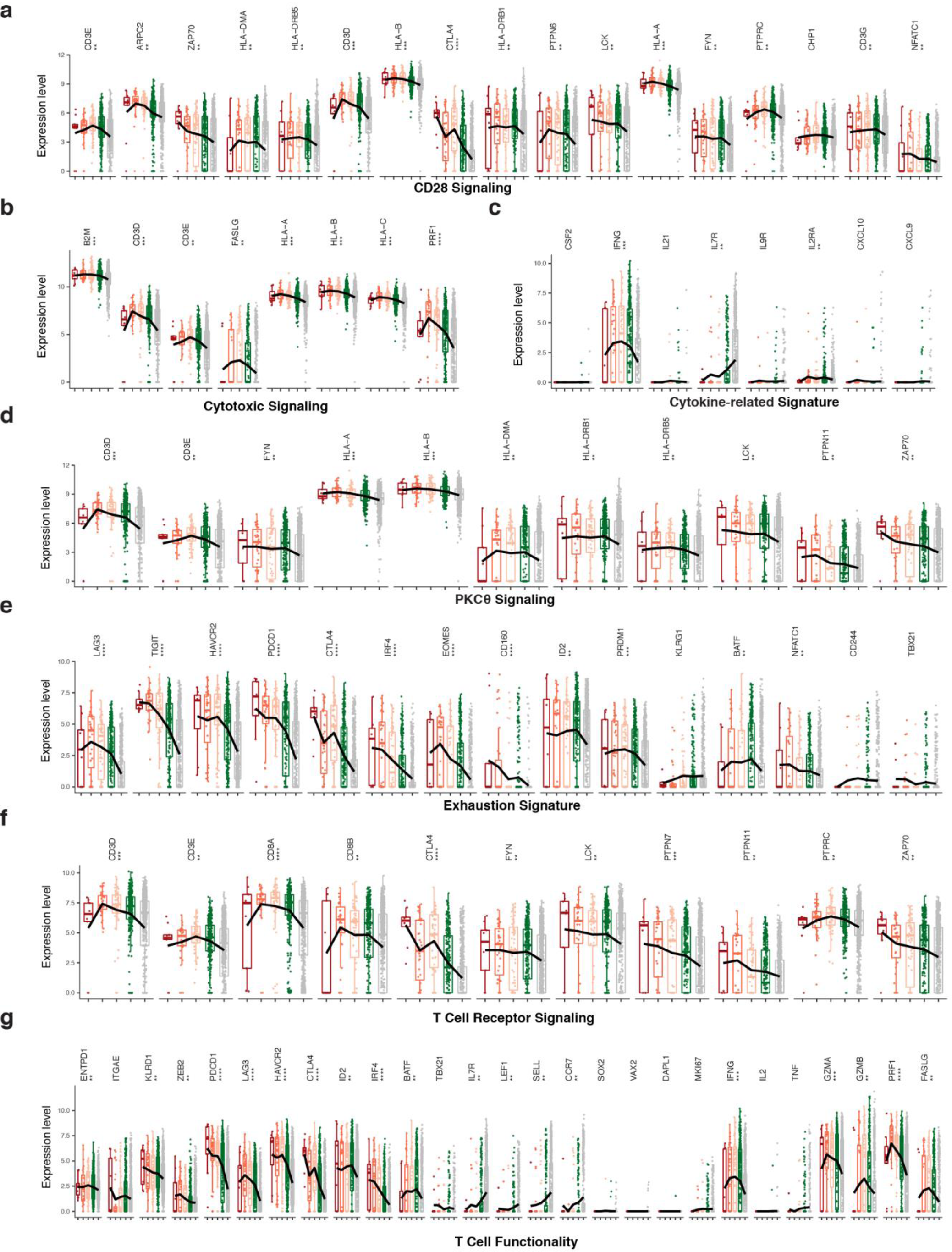
The statistical p values for the genes with differential expression among the TIL categories in Figs. 2c-d. The molecules of signaling pathways were derived from Ingenuity Pathway Analysis (IPA) by differential expression genes (Supplementary Table xx), **a,b,d,f**. And the gene signatures for **c,e,g** were derived from literature (**Supplementary Text**). The statistical significance was derived from One-way ANOVA analysis. * < 0.05, **<10e−4, ***<10e−8, ****<10e−16.

**Extended Data Figure 8.**
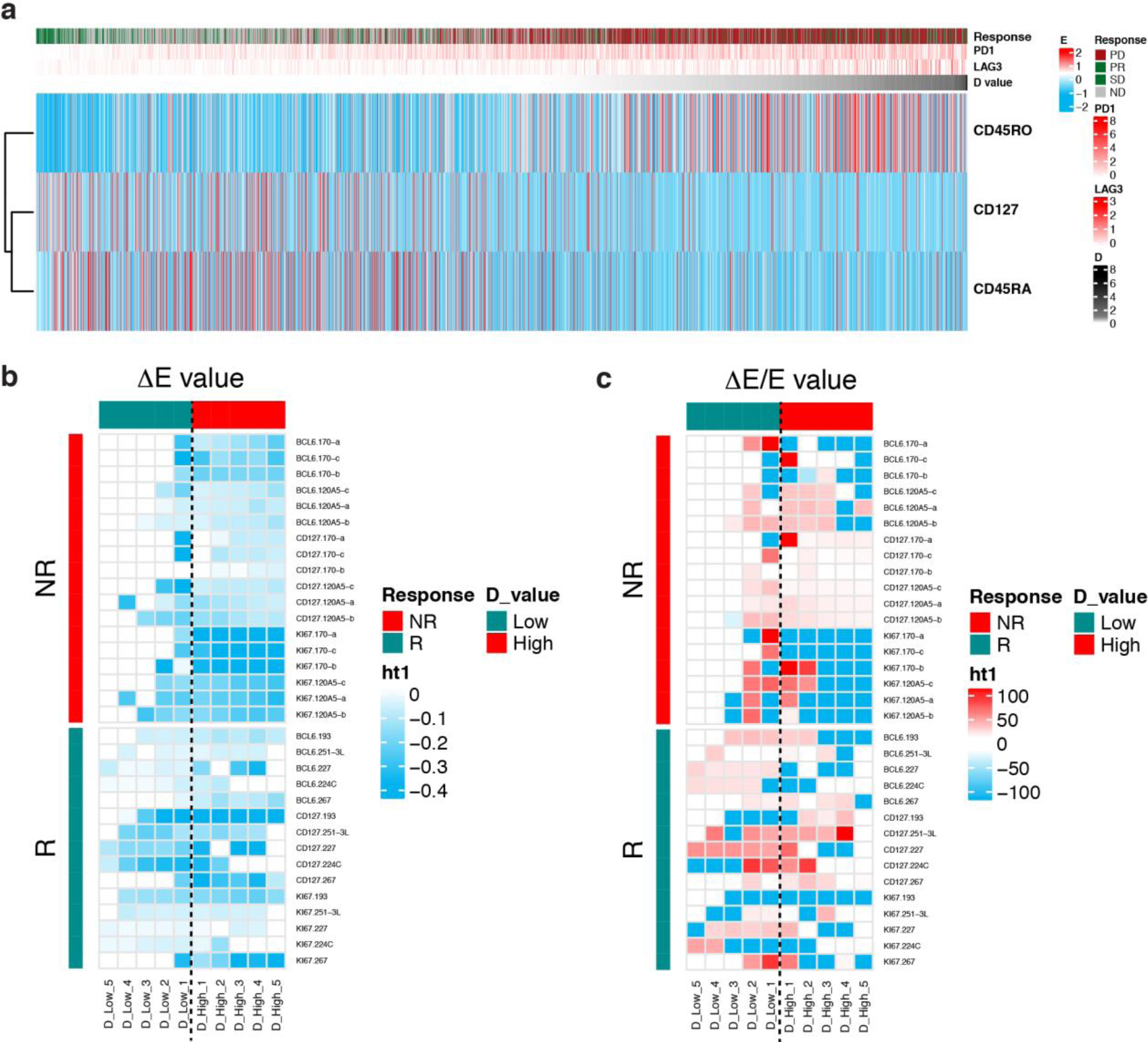
Effector and memory TILs sorted by D values of PF TILs from the human CyTOF data and the heatmaps for another 3 negative markers for exhaustion. **(a)** Hierarchical clustering of the effector and memory markers, including CD45RO, CD45RA, and CD127 by the expression values of these markers. **(b)** The ΔE values for the negative exhaustion markers, BCL6, CD127, and Ki-67, Supplementary to **Fig. 1g**. **(c)** The ΔE/E values for the negative exhaustion markers, BCL6, CD127, and Ki-67, Supplementary to **Fig. 3c**.

**Extended Data Figure 9.**
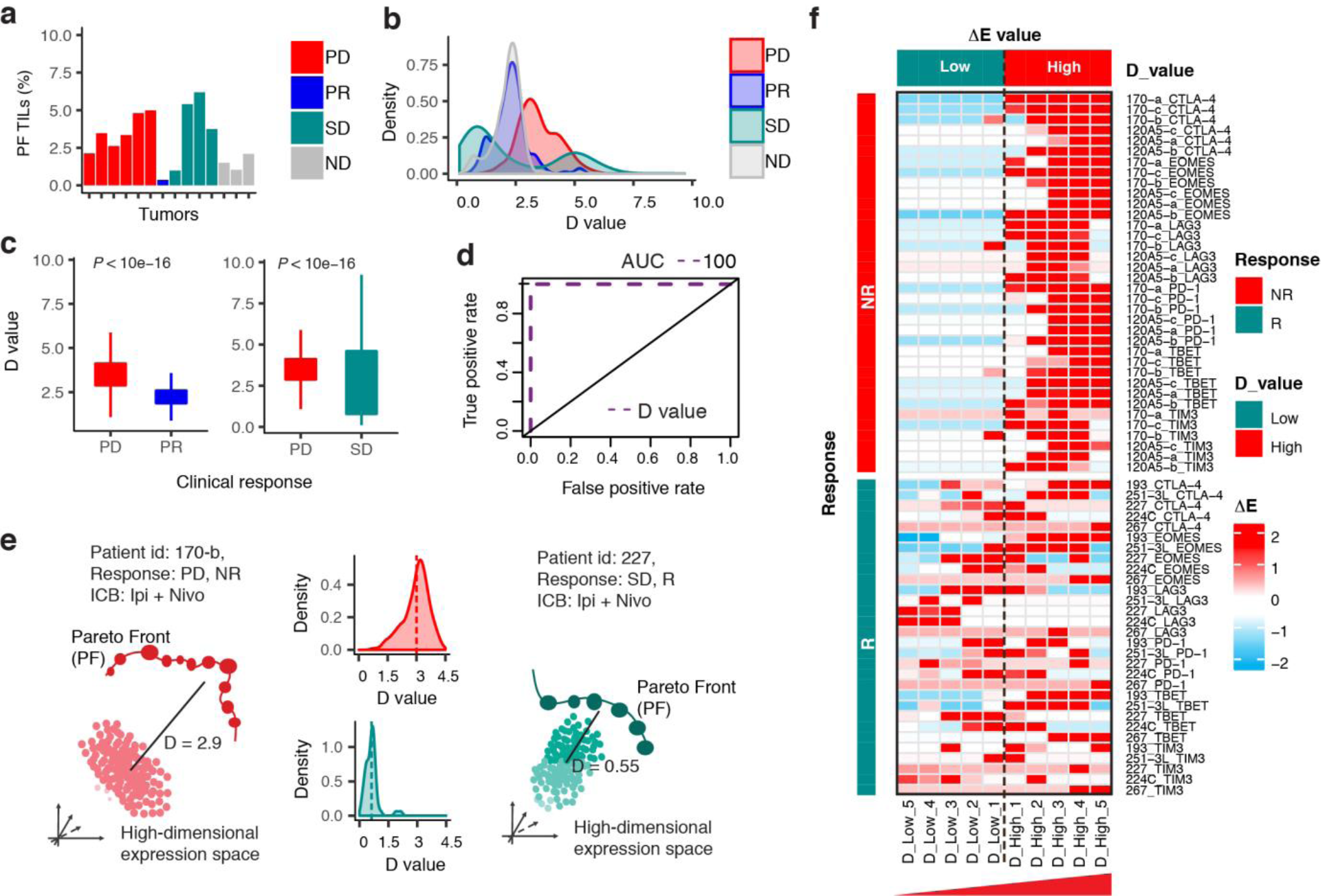
D value identified from 6 exhaustion markers by HD-scMed in distinguishing clinical responses to ICB. Clinical responses include PD: progressive disease, PR: partial response, and SD: stable disease. ICBs are αPD-1, αCTLA-4, and αPD-1+αCTLA-4. ND: PBMC samples from healthy donors were used as the baseline for ICB samples. **(a).** Identification of **Pareto front (PF)** TILs as exhausted TILs in the high-dimensional expression space. TIL exhaustion is evaluated by HD-scMed in the high-dimensional expression space defined by αPD-1, CTLA4, LAG3, EOMES, TIM3, and TBET, from human CyTOF antibody panel (**Online Methods**, **Extended Data Fig. 1**). Shown is a small portion of PF TILs identified as exhausted TILs by HD-scMed. **(b).** D value distributions in tumors with different clinical responses. D value is defined as the Euclidean distance or straight-line distance in the 6-dimensional expression space by the expression levels of the 6 exhaustion markers. D value for each PF TIL is the Euclidean distance between the PF TIL and its baseline non-front TILs that are dominated by the PF TIL (**Extended Data Fig.1a**). **(c).** Analysis of D values for PF TILs of the tumors from non-responders (PD) versus responders (PR or SD). P < 10e−16, Mann-Whitney *U* test. **(d).** ROC curve. The prediction accuracy of D value in distinguishing the tumors of non-responders from those of responders is AUC =100. 5-fold cross-validation supporting vector machine (SVM). AUC: The area under the ROC curve. ROC: Receiver Operating Characteristic. **(e)**. An instance to show the difference of D-value in the TILs of a non-responder (NR) and a responder (R). Both patients received the combination therapy of αPD-1+αCTLA-4 by Ipi+Nivo. Tumor 170-b is from the NR patient with progressive disease whereas tumor 227 is from the R patient with stable disease. The D-value for each tumor is averaged from those of PF TILs of this tumor. **(f).** Heatmap of ΔE values of the 6 exhaustion markers by the TILs from D low to D high. 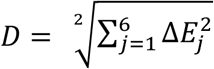, where ΔE value of each marker is the expression level difference between the PF TILs and their baseline non-front TILs. D value category is classified by D values of the PF TILs.

**Extended Data Figure 10.**
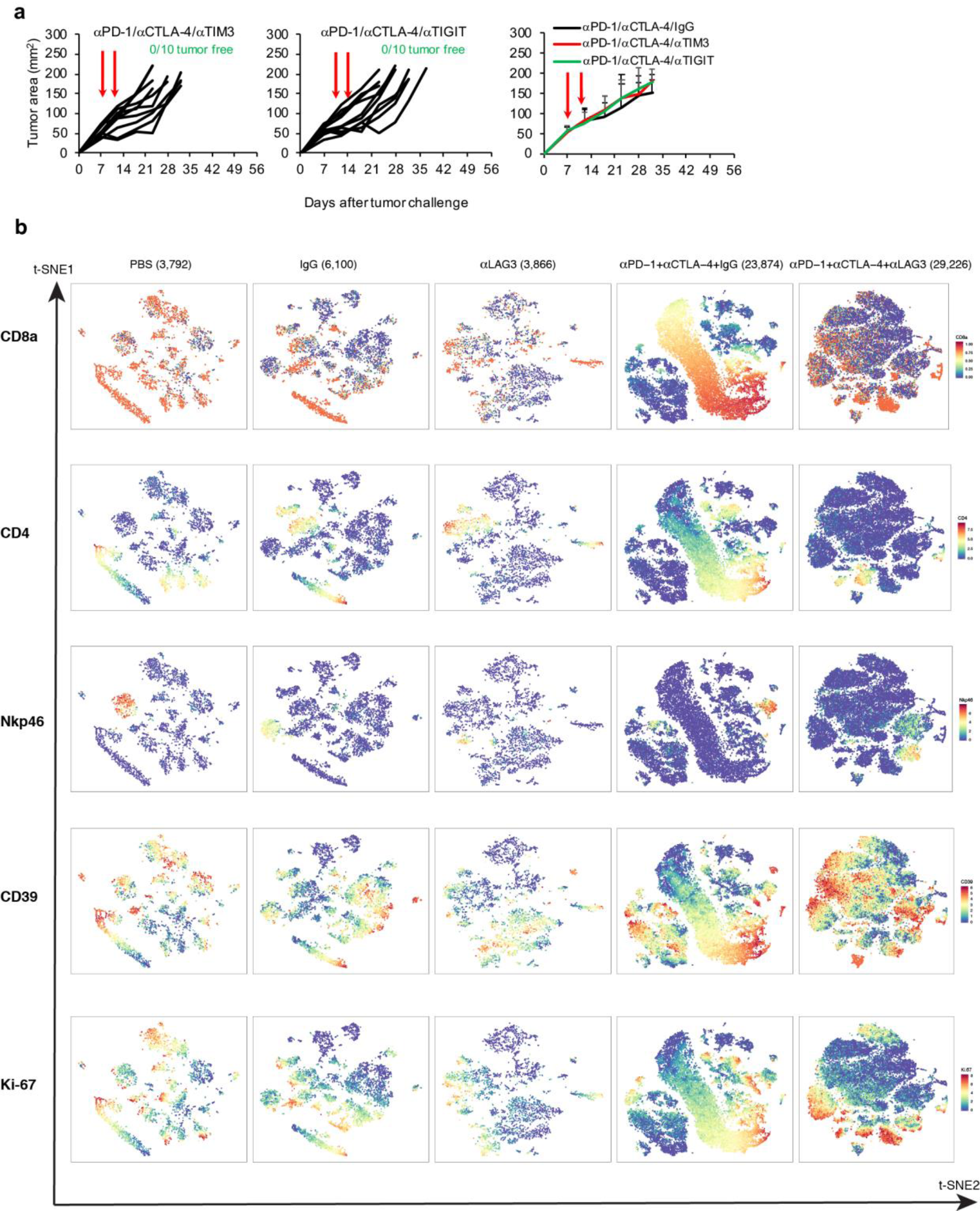
In vivo experiments for αPD-1+αCTLA-4+αTIM3 and αPD-1+αCTLA-4+α TIGIT and the expression levels of the markers from CD26 mouse CyTOF data. **(a)** CT26 tumor growth in BALB/c mice that were treated αPD-1+αCTLA-4+αTIM3 or αPD-1+αCTLA-4+αTIGIT or αPD-1+αCTLA-4+IgG as indicated by arrow (n=10). (**b**) t=SNE plot for the markers: CD8a, CD4, Nkp46, CD39, Ki-67.

